# A targeted amplicon sequencing panel for cost-effective high-throughput genotyping of *Aedes aegypti*

**DOI:** 10.1101/2025.09.28.679043

**Authors:** Josquin Daron, Alicia Lecuyer, Laurence Ma, Pegah Marzooghi, Mallery I. Breban, Peter Kyrylos, I’ah Donovan-Banfield, Seth N. Redmond, Louis Lambrechts

## Abstract

The mosquito *Aedes aegypti* is the primary vector for several medically significant arboviruses, including dengue, Zika, chikungunya, and yellow fever. Studying the genetic diversity of *Ae. aegypti* is crucial for understanding its evolutionary history, population dynamics, and the effectiveness of vector control methods. Due to the large genome size of *Ae. aegypti*, whole-genome sequencing (WGS) is often cost-prohibitive for large-scale studies. Recent advances in reduced-representation methods, aiming to reduce costs by sequencing only a small portion of the genome, provide cost-effective alternatives. However, a standardized set of genome-wide markers specifically designed for population genetic studies of *Ae. aegypti* remains unavailable. Here, we present a targeted amplicon sequencing panel designed for cost-effective, high-throughput genotyping across 291 loci distributed throughout the *Ae. aegypti* genome. Our *in silico* analyses indicate that this amplicon panel effectively replicates population structure analyses typically derived from WGS data. We demonstrate that the amplicon panel accurately discriminates between diverse laboratory colonies of *Ae. aegypti* and consistently measures individual genetic admixture to a degree comparable with WGS. By enabling high-throughput genotyping at a reduced cost, we anticipate that our targeted amplicon sequencing panel will facilitate large-scale genotyping studies of *Ae. aegypti* for vector surveillance and population structure analyses, especially in resource-limited settings.

## Introduction

The mosquito *Aedes aegypti* is the primary vector of multiple arthropod-borne viruses (arboviruses) of medical significance such as dengue, Zika, chikungunya and yellow fever (Powell et al., 2018). Today, *Ae. aegypti* is prevalent throughout the tropics and subtropics, and its range is projected to expand further due to urbanization, connectivity and climate change (Kraemer et al., 2019). The species consists of two subspecies: *Aedes aegypti formosus* (*Aaf*), a generalist found in both forests and urban areas within its native African range, and the globally invasive *Aedes aegypti aegypti* (*Aaa*), a human specialist that thrives in urban environments (Gloria-Soria et al., 2016; Powell and Tabachnick, 2013). The human specialist *Aaa* evolved from generalist ancestors in West Africa approximately 5,000 years ago (Rose et al., 2023), facilitating its global expansion over recent centuries. Its preference for human hosts, coupled with its higher vector competence, renders *Aaa* a highly effective transmitter of human arboviruses (Aubry et al., 2020; Caldwell et al., 2024; Rose et al., 2022; Rose et al., 2020). The dichotomy between *Aaa* and *Aaf* breaks down in some African locations where *Ae. aegypti* populations display patterns of genetic admixture between *Aaa* and *Aaf*, reflecting mixed genetic ancestry (Crawford et al., 2025; Kotsakiozi et al., 2018a; Lozada-Chavez et al., 2025; Rose et al., 2022; Rose et al., 2020). There is also extensive genetic diversity within each *Ae. aegypti* subspecies, particularly within *Aaf* (Crawford et al., 2025; Lozada-Chavez et al., 2025).

Population genetic studies of *Ae. aegypti* provide insights into the evolutionary pathways and genetic variations that have shaped this mosquito species over time. By examining its genetic structure, we can better understand how different environmental and biological factors influence population size, gene flow, and the spread of adaptive traits (Crawford et al., 2025; Lozada-Chavez et al., 2025; Rose et al., 2023; Rose et al., 2020). By genetically monitoring mosquito populations, we can also examine the direct effect of vector control interventions by tracking changes in vector population size and dispersal rates (Filipovic et al., 2020; Jasper et al., 2019).

In the pre-genomic era, population genetic studies of *Ae. aegypti* mainly relied on small sets of genetic markers such as microsatellites. These simple sequence repeats are highly informative but their number in the *Ae. aegypti* genome is limited (Brown et al., 2011; Lovin et al., 2009). Moreover, they are prone to human error during scoring and require careful consideration for cross-study comparisons, complicating their use as high-throughput markers (Pasqualotto et al., 2007; Van Oosterhout et al., 2004). With recent advances in high-throughput sequencing technologies, single-nucleotide polymorphisms (SNPs) and genotyping-by-sequencing methods have gained traction (Davey et al., 2011). Unlike microsatellites, SNPs are abundant and straightforward, though they provide less information per marker as they are typically biallelic. However, their abundance allows large-scale screening to offset this limitation (Smouse, 2010). Despite declining costs of high-throughput sequencing technologies, whole-genome sequencing (WGS) remains cost-prohibitive for large-scale genotyping studies of *Ae. aegypti* due to its large genome size (1.3 Gb) and high repeat content (>60%) (Daron et al., 2025; Daron et al., 2024; Matthews et al., 2018).

In the last two decades, several alternative SNP-based genotyping methods to WGS have been developed, aiming to reduce costs by sequencing only a small portion of the genome, a concept known as reduced-representation approaches (Davey et al., 2011). Restriction site-associated DNA sequencing (RAD-seq) has become a popular approach for identifying and screening SNPs (Peterson et al., 2012) and has been successfully applied to *Ae. aegypti* (Fontaine et al., 2017; Rasic et al., 2014; Rasic et al., 2015; Schmidt et al., 2018). Despite its usefulness, RAD-seq poses analytical challenges due to pervasive missing data (Arnold et al., 2013; Huang and Knowles, 2016). Building on RAD-seq data, a high-throughput genotyping chip with over 25,000 validated SNPs was developed for *Ae. aegypti* (Evans et al., 2015) and effectively used in studies of population structure and demographic inference (Kotsakiozi et al., 2018a; Kotsakiozi et al., 2018b; Soghigian et al., 2020). This genotyping chip offers a more accurate and cost-effective alternative to low-coverage WGS (<10×) for population genetic studies (Gomez-Palacio et al., 2024). However, its reliance on a pre-selected set of SNPs introduces ascertainment bias (Albrechtsen et al., 2010). Alternatively, low-coverage WGS has recently emerged as a cost-effective approach for genome-wide screening on a population scale, offering costs comparable to RAD-seq while providing comprehensive genomic data (Lou et al., 2021). Nevertheless, this method presents a trade-off by reducing coverage depth, which reduces confidence in individual genotype calls, and still entails relatively high library preparation costs and bioinformatic workload.

Targeted amplicon sequencing panels offer another valuable reduced-representation strategy for conducting cost-effective genotyping studies on a large scale. These panels are designed to simultaneously amplify specific genomic loci using polymerase chain reaction (PCR). Barcoding of individual samples allows subsequent high-throughput sequencing of a multiplexed library. The resulting sequences are then compared with a reference genome to call SNPs and conduct various analyses. Furthermore, by focusing on sequenced regions containing multiple SNPs, one can determine “microhaplotypes” using phased short-read sequences (Baetscher et al., 2018). Multi-allelic marker data from these loci significantly increase the power for relationship inference, such as estimating identity by descent (Siegel et al., 2024). These targeted amplicon panels have been developed for various applications, including vector species identification and pathogen detection (Batovska et al., 2018; Cannon et al., 2021; Lima-Cordon et al., 2025; Makunin et al., 2022), as well as for screening insecticide resistance (Campos et al., 2022; Collins et al., 2022; Fontaine et al., 2024; Lima-Cordon et al., 2025). However, no targeted amplicon sequencing panel is currently available for population genetic studies of *Ae. aegypti*.

Here, we introduce a targeted amplicon sequencing approach that genotypes 291 loci across the *Ae. aegypti* genome. We provide evidence that this amplicon panel provides a cost-effective alternative to WGS for population genetic analyses, supporting its use for large-scale genotyping studies in resource-limited settings.

## Materials and methods

### Amplicon panel design

Target loci were selected based on the AaegL5 assembly (Matthews et al., 2018) and the Aaeg1200 genomes sequencing dataset (Crawford et al., 2025). Unless otherwise stated, all statistics were calculated using scikit-allel v1.3.1 (Miles et al., 2024). Mean pairwise nucleotide diversity (π) was called in 50-base pair (bp) windows across the genome, and candidate target regions were identified as 50-to 150-bp high diversity regions (π>0.005) flanked by low diversity regions (π<0.0001) to design PCR primers. Regions with low diversity and known transposable elements identified in the AaegL5 assembly (Matthews et al., 2018) were removed.

The remaining ∼32,000 candidate regions were filtered to remove sites where high sequencing depth (>20,000× in all samples) and/or low diversity (π<0.02, <25 SNPs per locus) might indicate a potential copy-number polymorphism. Low-depth samples (<5,000×) were removed to ensure high call rates. Target regions exhibiting strong selection signals (–2<Tajima’s D<2) (Tajima, 1989) or inbreeding (Malecot’s *f*<0.25) (Malécot, 1948) in either *Aaa* or *Aaf* populations were also removed. Out of 16,557 target regions that passed all filters, 10 sets of 350 potential loci each were pseudo-randomly selected using a gap-filling algorithm to optimize for both SNP and centimorgan (cM) distances (Juneja et al., 2014).

Primer design was carried out at GT-seek LLC (https://gtseek.com) for primers with a suitable melting temperature and target size to perform multiplex amplification (Campbell et al., 2015). Primers were designed to include a locus-specific sequence of 17-26 bp and an Illumina sequencing adapter of 31 bp (forward) or 34 bp (reverse). Potential dimer pairs were removed, and a set of 300 target loci was chosen to optimize for cM gap length. During optimization of the sequencing library preparation, 9 primer pairs were subsequently excluded due to amplification artefacts. The final 291-loci panel had a median distance of 0.3 cM between markers. All primer sequences are provided in Table S1.

### DNA extraction

DNA was extracted from individual mosquitoes with DNeasy 96 Blood & Tissue Kit (Qiagen), DNAzol DIRECT (Molecular Research Center Inc.), or the Pat-Roman protocol as previously described (Dickson et al., 2020). Both commercial kits were used following their manufacturer’s instructions. The Pat-Roman protocol is an inexpensive method relying on a homemade buffer to lyse the tissues followed by a series of mixing, incubating, and centrifugation steps to isolate the DNA. Briefly, the mosquito was homogenized in a mixer mill (Precellys 24, Bertin Technologies) in Pat Roman’s buffer (0.1 M NaCl, 0.2 M sucrose, 0.1 M Tris-Hcl pH 8.0, 0.05 M EDTA pH 8.0, 0.5% SDS, adjusted at pH 9.2). The lysate was incubated at 65°C and 8 M potassium acetate was added. After incubation on ice, the lysate was transferred to a new tube and centrifugated. Supernatant was transferred to a fresh tube and mixed with 96% ethanol to precipitate DNA overnight at –20°C. DNA was centrifugated and the pellet was washed with ice-cold ethanol and resuspended in TE buffer (Fisher Scientific).

### Library preparation and sequencing

Amplicons were generated in two successive PCRs (Figure 1). In the first PCR (PCR1), all 291 target loci were amplified simultaneously in a multiplex reaction, using a balanced primer pool. The PCR1 primers were initially resuspended in nuclease-free water to a concentration of 200 μM. To remove extremes of sequencing coverage, each primer was then diluted individually according to a balancing factor based on initial test sequencing, as detailed in Table S1. Equal volumes of each diluted primer were combined in a single tube to create the balanced primer pool, resulting in an average concentration of ∼50 nM per primer.

**Figure 1:**
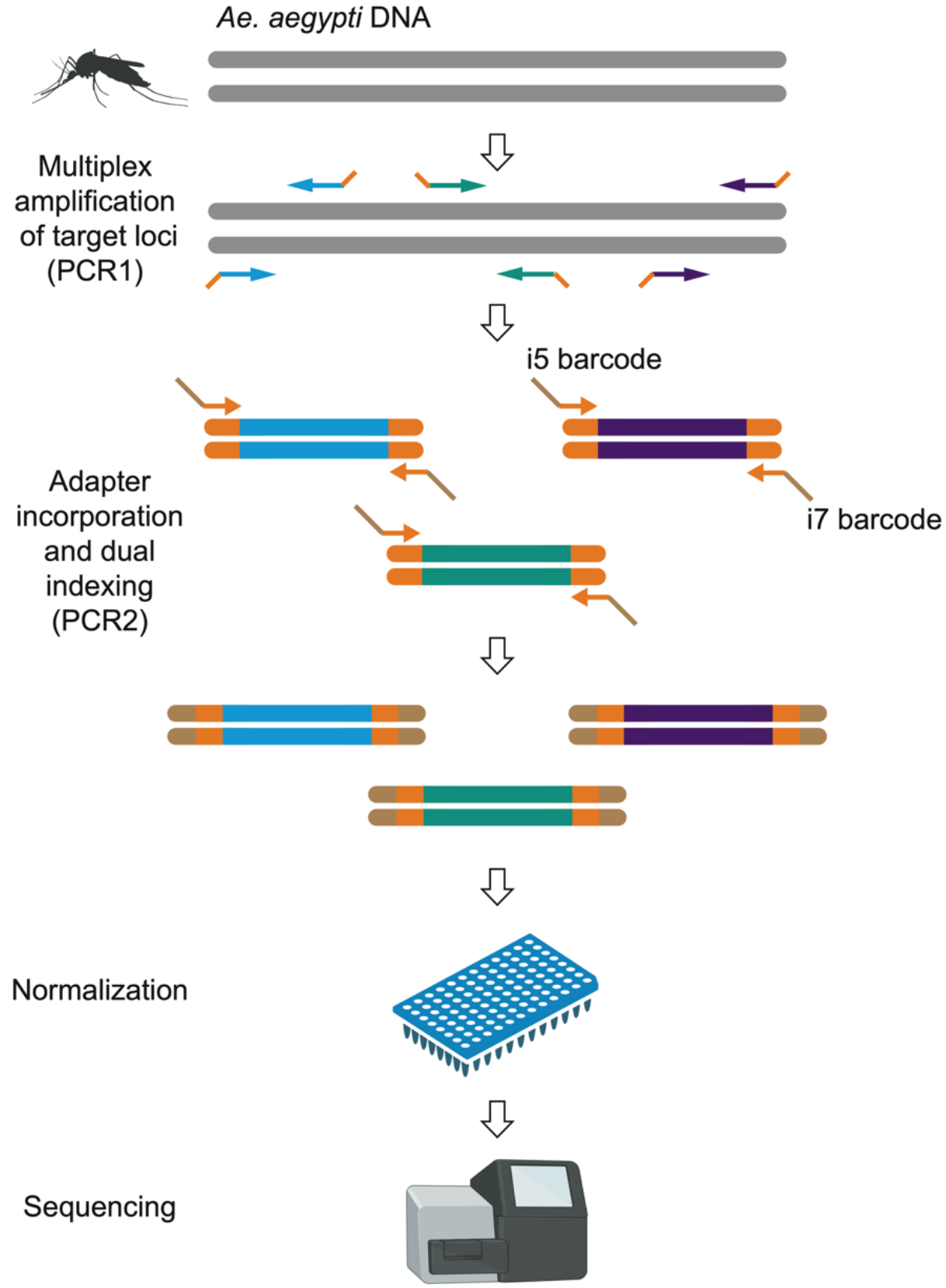
Targeted amplicon sequencing workflow. Amplicons were generated through two rounds of PCR. The first PCR (PCR1) involved amplifying 291 target loci simultaneously using a balanced primer pool, followed by a second PCR (PCR2) during which Illumina capture sites and unique dual indexes were added. The final products were normalized, pooled, and sequenced on an Illumina MiSeq platform.

All PCR1 components were thawed at room temperature (20-25°C), mixed well, and briefly centrifuged. For each sample, 2 μl of genomic DNA was mixed with 3.5 μl of Qiagen Plus Multiplex Master Mix 2× and 3 μl of the balanced primer pool in a 96-well PCR plate (Fisher Scientific). The PCR1 cycling conditions were as follows. The reaction was initiated with a hot start at 95°C for 15 min. A denaturation step was then conducted at 95°C for 30 sec, followed by a gradual ramp (5% or 0.2°C/sec) to 57°C for a 2-min annealing step, and an extension step at 72°C for 30 sec, repeated for five cycles. This was followed by 15 cycles of denaturation at 95°C for 30 sec, annealing at 65°C for 30 sec, and extension at 72°C for 30 sec.

In the second PCR (PCR2), Illumina capture sites (P5 and P7) and unique dual indexes (i5 and i7) were appended to the PCR1 products by using PCR2 primers paired for combinatorial indexing (Table S1). The PCR1 products were diluted 20-fold in nuclease-free water and 3 μl of the diluted PCR1 products were mixed in a 96-well PCR plate with 5 μl of Qiagen Plus Multiplex Master Mix 2× and 1 μl of each PCR2 primer diluted at 2.5 μM. The PCR2 cycling conditions were as follows. The reaction was initiated with a hot start at 95°C for 15 min. A denaturation step was then conducted at 95°C for 10 sec, followed by an annealing step at 65°C for 30 sec, and an extension step at 72°C for 30 sec, repeated for 15 cycles. A final extension was performed at 72°C for 5 min.

PCR2 products were normalized using either a SequalPrep Normalization Plate Kit 96-well (Fisher Scientific), according to the manufacturer’s instructions, or manually to match the lowest concentration sample, using a Qubit dsDNA HS Assay Kit (Invitrogen). Each normalized sample contributed 10 μl to a combined pool in a single tube. From this pool, 50 μl was mixed in a new tube with 40 μl of AMPure XP magnetic beads (Agencourt) at a ratio of 0.8× and incubated at room temperature for 10 min. After placing the tube on a magnetic rack for another 10 min to let the solution resolve, the cleared supernatant was removed and discarded. While still on the rack, 200 μl of freshly-prepared 70% ethanol was added, incubated at room temperature for 30 sec, and then discarded; this wash step was repeated once. After the second wash, the tube was taken off the rack and left open for 2 min at room temperature to slightly desiccate the beads. To elute the purified product, 15 μl of TE buffer (Fisher Scientific) was added to the dried beads, thoroughly mixed, and returned to the magnetic rack. Once the solution cleared, the supernatant containing the purified library pool was collected and transferred to a new tube. The expected library size (218-305 bp) was verified using Bioanalyzer High Sensitivity DNA Kit (Agilent). Before purification, primer-adapter dimers often appeared as a secondary peak of ∼170 bp, but they were generally eliminated by purification. If needed, additional removal could be achieved by performing a gel clean-up and/or slightly reducing primer concentration. Sequencing was performed on a MiSeq platform (Illumina) using paired-end 2×150-bp reads, with either a MiSeq Reagent v2 Micro flowcell (yielding approximately 8 million reads) or MiSeq Reagent v2 Nano flowcell (yielding approximately 2 million reads).

### Processing of amplicon sequencing reads

The sequencing reads were initially demultiplexed by sample using the dual-index barcodes on the Illumina platform, resulting in one FASTQ file per sample. The raw reads were trimmed using Cutadapt v2.10 (parameters: -q 30 -m 50 --max-n 0) (Martin, 2011) and mapped to the AaegL5 reference genome assembly (Matthews et al., 2018) using BWA-MEM v0.7.17 (Li, 2013) with default parameters. PCR duplicates were removed using Picard Tools (http://broadinstitute.github.io/picard), and SNPs were called using GATK HaplotypeCaller v4.1.9.0 (McKenna et al., 2010). Only SNP variants were selected, and low-quality SNPs were excluded if they failed any of the following criteria: QD>5, FS<60, and ReadPosRankSum>–8. Genotypes with a genotype quality (GQ) ≥30 and sequencing depth ≥10× were retained. To minimize missing data in the final SNP matrix, variants and individuals with >10% missing data were discarded.

### Processing of WGS reads

The same genotyping procedure as for amplicon sequencing reads was applied to 656 publicly available WGS datasets for *Ae. aegypti* retrieved from NCBI bioprojects PRJNA602495 (Rose et al., 2020), PRJNA385349 (Kelly et al., 2021), PRJNA882905 (Rose et al., 2022), PRJNA864744 (Love et al., 2023), and PRJNA943178 (Lozada-Chavez et al., 2025). The processing of these datasets followed the approach described above, with three adjustments made to account for the heterogeneous sequencing depth. First, the GQ threshold was relaxed to include genotypes with a GQ>20 and a sequencing depth ≥5×. Second, variants with >10% missing data and individuals with >50% missing data were discarded. Third, the degree of relatedness among individuals (kinship coefficient) was estimated with plink v1.9 (parameter: -- genome) (Purcell et al., 2007) and samples showing a low level of relatedness (PI_HAT>0.75) were retained for subsequent population genetic analyses. This process ultimately yielded a final matrix of 493 individuals × 15 million SNPs.

### Putative off-target loci

To identify putative off-targets of the amplicon panel, each PCR1 primer pair was aligned against the reference genome using the primersearch tool from the EMBOSS suite (Rice et al., 2000), with the option -mismatchpercent 5, allowing for a single mismatch. The number of times each primer pair was mapped to the genome within a maximum distance of 300 bp was then calculated. Primer pairs producing more than one match in the genome were considered to have putative off-target loci.

### Population genetics

To explore population structure, SNPs in linkage disequilibrium were first removed by excluding those with an *r*^*2*^>0.01 within moving windows of 500 SNPs and a step size of 250 SNPs, using scikit-allel v1.3.3 (Miles et al., 2024). The population genetic structure was visualized using principal component analysis (PCA) in scikit-allel. Individual genetic ancestry proportions were quantified using ADMIXTURE v1.3.0 (Alexander et al., 2009), with the number of clusters (K) tested from 2 to 5. For the WGS dataset, ADMIXTURE was run 100 times for each K value, with each dataset comprising a random sampling of 100,000 unlinked variants. The most likely number of ancestral populations (K) was determined using the cross-validation (CV) error rate (Alexander et al., 2009). The lowest CV error rate indicated that K=2 ancestral populations was the most likely model.

## Results

### Genetic diversity of natural *Ae. aegypti* populations at amplicon loci

We designed an amplicon panel of 291 loci collectively representing 41,423 bp (0.003% of the total *Ae. aegypti* genome size). The size of amplicons ranges from 87 bp to 174 bp with a median of 143 bp (Figure 2A). We first aimed to determine if the genetic diversity captured by this amplicon panel recapitulates the population structure and diversity of *Ae. aegypti* observed on genome-wide scale. We conducted a series of analyses based on the amplicon sequences extracted from publicly available WGS data for *Ae. aegypti* (Kelly et al., 2021; Love et al., 2023; Lozada-Chavez et al., 2025; Rose et al., 2022; Rose et al., 2020). From approximately 15 million SNPs identified, we extracted a set of 2,826 SNPs across 493 individuals, present exclusively at the amplicon loci. These SNPs were evenly distributed across the genome, with a median of 12 SNPs per amplicon (Figure 2B). We found that chromosomes 2 and 3 were particularly well covered, whereas the first 100 Mb of chromosome 1 exhibited the lowest density of suitable sites and SNPs. Our analysis revealed a higher number of SNPs in African *Ae. aegypti* populations compared to those outside of Africa (Figure 2C), consistent with earlier population genomic studies reporting greater genetic diversity in African populations (Crawford et al., 2025; Lozada-Chavez et al., 2025).

**Figure 2:**
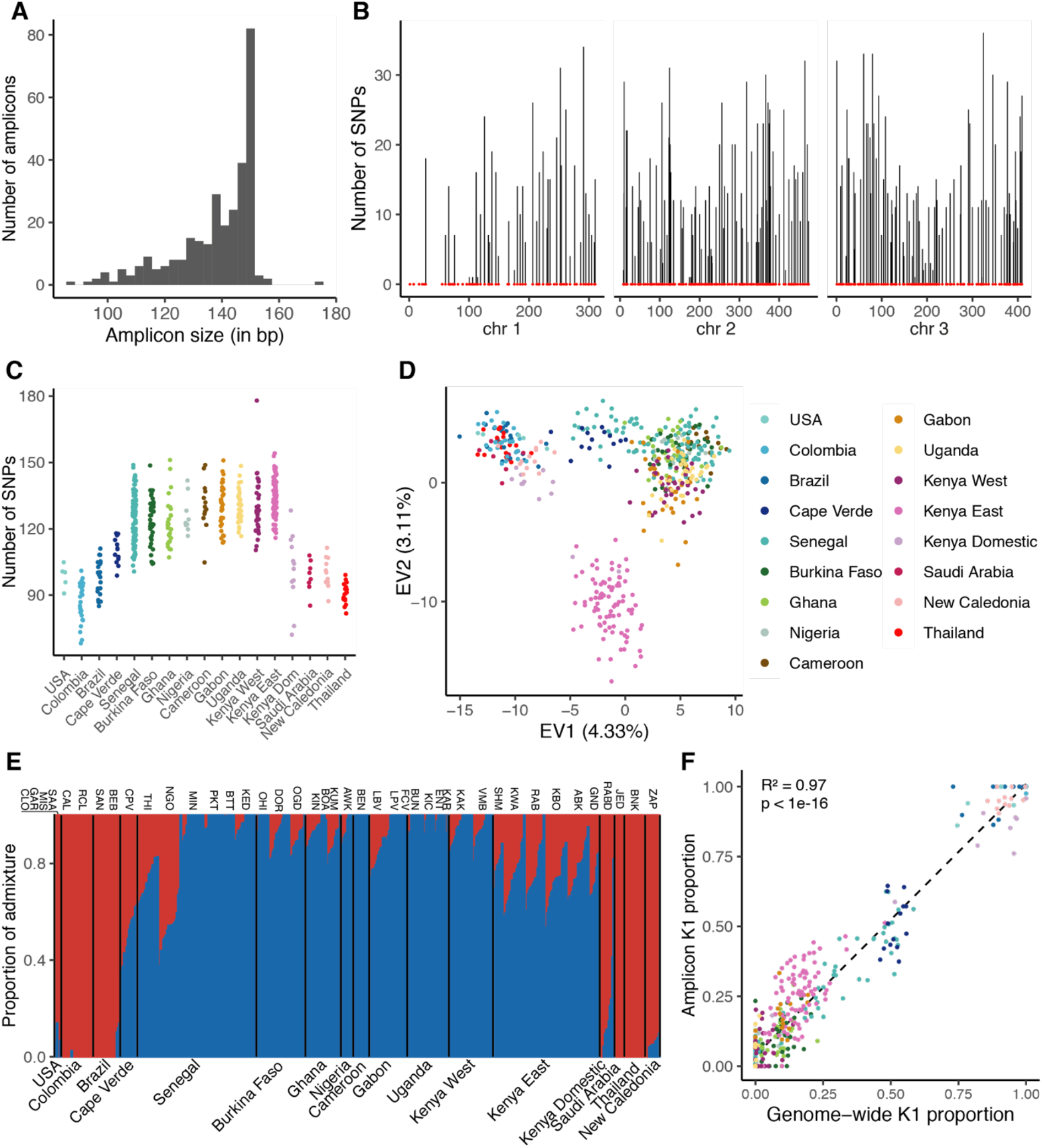
Global genetic diversity and population structure of natural *Ae. Aegypti* populations inferred from amplicon loci. (**A**) Predicted amplicon size distribution. (**B**) Genomic distribution of amplicons (red dots) and number of SNPs identified at amplicon loci (vertical bars) along the three chromosomes (chr, shown in Mb) in natural populations worldwide (chr=chromosome). (**C**) Total number of SNPs identified per individual. (**D**) PCA of global genetic diversity at amplicon loci (EV=eigenvalue). (**E**) Barplot of admixture proportions per individual assuming two ancestry components (K=2, with *Aaa* ancestry in red and *Aaf* ancestry in blue). (**F**) Correlation between the proportion of *Aaa* ancestry (K1) estimated from genome-wide data and from amplicon loci.

We next investigated the genetic structure of global populations using a set of unlinked SNPs. Principal component analysis (PCA) broadly mirrored patterns observed at the whole-genome level (Crawford et al., 2025; Lozada-Chavez et al., 2025; Rose et al., 2020), revealing three distinct clusters (Figure 2D). African populations were separated into two clusters: one comprising all populations from West of the Rift Valley, and another with two Kenyan populations from East of the Rift Valley. The third cluster consisted of populations from outside Africa, representing the invasive *Aaa* subspecies that expanded globally in the last few centuries (Rose et al., 2023). Although this pattern matches what was observed in previous studies (Crawford et al., 2025; Lozada-Chavez et al., 2025), the resolution was lower with amplicon loci compared to genome-wide markers, likely due to fewer variants and shallow WGS coverage (mean 13.59×, min 0.55×, max 35×).

Using Bayesian clustering, we characterized the genetic ancestry of the global samples, identifying two main ancestry components corresponding to *Aaa* (K1) and *Aaf* (K2) (Figure 2E). As expected, several *Ae. aegypti* populations showed significant levels of admixture between K1 and K2. The proportion of K2 ancestry estimated from the amplicon loci was strongly correlated with the proportion estimated from WGS data (R^2^=0.97, p<1×10−^16^; Figure 2F), demonstrating the ability of the amplicon panel to infer ancestry proportions.

### Genetic admixture in Cape Verde *Ae. aegypti*

Building on these results, we assessed the accuracy of the amplicon panel to infer mixed genetic ancestry. We genotyped 15 wild-caught *Ae. aegypti* samples from Cape Verde using the amplicon sequencing strategy, chosen because earlier WGS highlighted significant admixture between *Aaf* and *Aaa* (23% *Aaa* ancestry on average) in these samples (Rose et al., 2022). Quality control showed successful mapping to amplicon loci for all but one sample, and a relatively even read coverage across the other 14 samples (Figure 3A). The distribution of read counts among amplicons was moderately skewed, with the most abundant amplicon accounting for ∼13% of the reads (Figure 3B).

**Figure 3:**
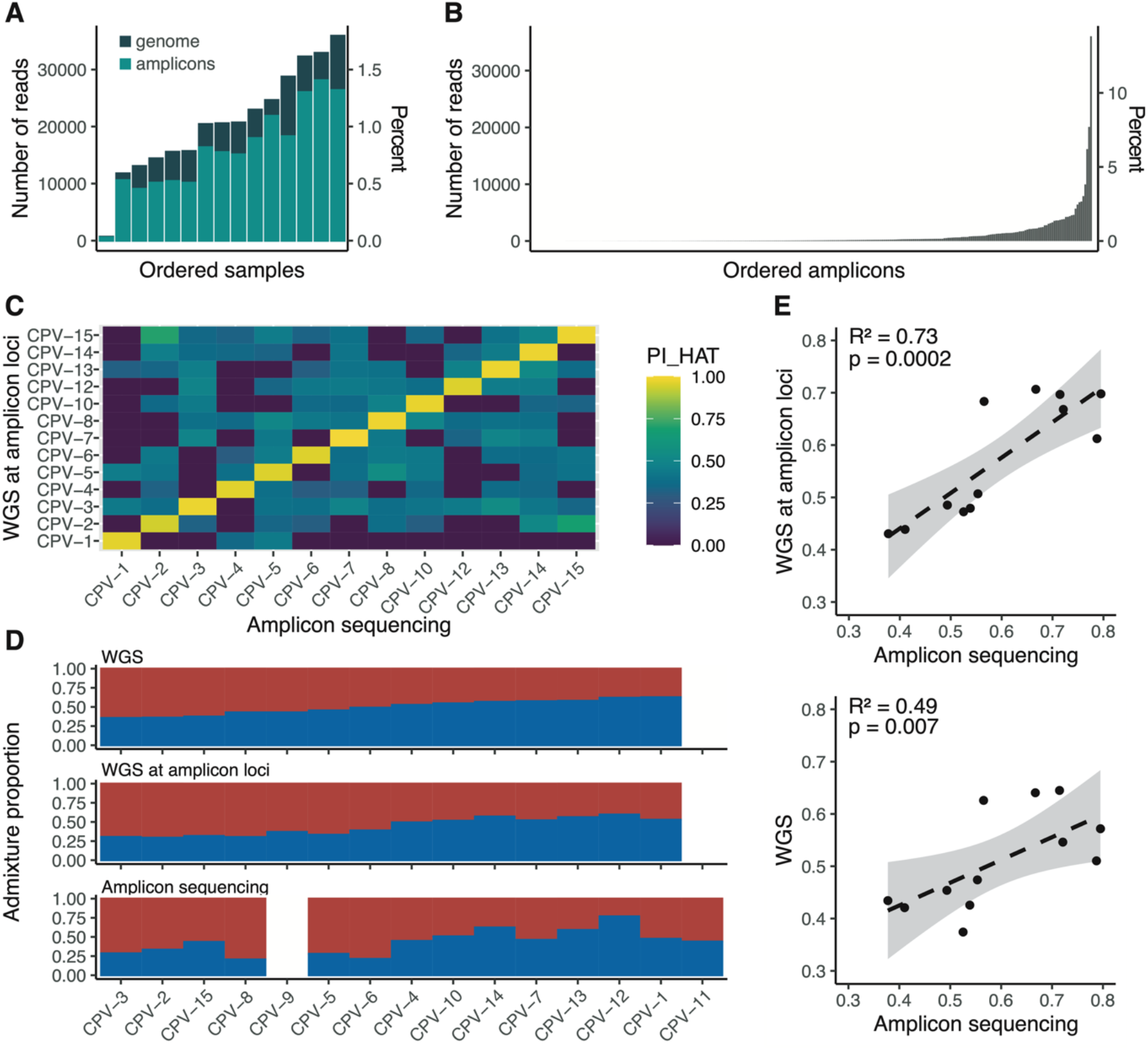
Admixture analysis of 15 wild-caught *Ae. aegypti* specimens from Cape Verde using amplicon sequencing vs. whole-genome sequencing. (**A**) Number of reads per Cape Verde (CPV) sample mapped to the amplicon loci (light green) or the AaegL5 assembly (dark green), respectively. (**B**) Number of reads uniquely mapping to each amplicon, ordered by increasing coverage. (**C**) Kinship (PI_HAT) matrix showing pairwise relatedness coefficients computed from amplicon loci genotyped using either WGS or amplicon sequencing. (**D**) Barplot of admixture proportions, assuming two ancestry components (K=2), estimated from WGS data (top), polymorphisms at amplicon loci genotyped by WGS (middle), and polymorphisms at amplicon loci genotyped by amplicon sequencing (bottom). (**E**) Correlation between the proportion of *Aaa* ancestry estimated at amplicon loci using WGS and amplicon sequencing (top), and between genome-wide polymorphisms and amplicon sequencing (bottom).

For the 14 individuals that passed quality control, we genotyped a total of 154 SNPs to evaluate genotype concordance between our amplicon sequencing data and WGS data through a relatedness analysis. Our pairwise comparisons showed high relatedness values between the same samples genotyped using the two different methods (Figure 3C).

We next investigated the genetic ancestry of the Cape Verde samples by analyzing three sources of polymorphism (Figure 3D): (i) genome-wide polymorphisms identified in WGS data, (ii) polymorphisms identified at amplicon loci extracted from WGS data, and (iii) polymorphisms identified through our amplicon sequencing approach. We found a strong positive correlation (R^2^=0.73, p=0.0002) between genetic ancestry proportions inferred from the amplicon loci using either amplicon sequencing or WGS data (Figure 3E). This correlation was slightly weaker but still statistically significant (R^2^=0.49, p=0.007) when comparing ancestry estimates from the amplicon loci to those derived from genome-wide polymorphisms (Figure 3E). These results demonstrate that the amplicon panel can accurately estimate the proportions of *Aaf* and *Aaa* ancestry components in admixed samples while using far fewer markers than WGS, thus validating the cost-effectiveness of the amplicon sequencing approach for genetic ancestry inference.

### Genetic diversity of laboratory *Ae. aegypti* colonies

To validate the proof of concept for our amplicon panel, we expanded the range of samples under investigation. We genotyped a total of 90 individuals from 12 different field-derived *Ae. aegypti* colonies after 7-29 generations of laboratory colonization. We generated between 2,352 and 137,943 reads per sample mapping to the amplicon loci (Figure 4A). The read coverage per amplicon was relatively uniform, with no amplicons disproportionately represented (Figure 4B). After removing five individuals presenting more than 10% missing data, a total of 1,373 SNPs were genotyped across the remaining 85 individuals. These SNPs were evenly distributed across the genome (Figure 4C). Consistent with earlier findings in natural populations of *Ae. aegypti* (Crawford et al., 2025; Lozada-Chavez et al., 2025), the number of SNPs per individual was higher in colonies originating from African populations than in those originating from non-African populations (Figure 4D). PCA revealed clear clustering of individuals within colonies (Figure 4E). Eigen value 1 (EV1) separated African from non-African colonies, while EV2 further differentiated African individuals along a West-to-Central-to-East African axis.

**Figure 4:**
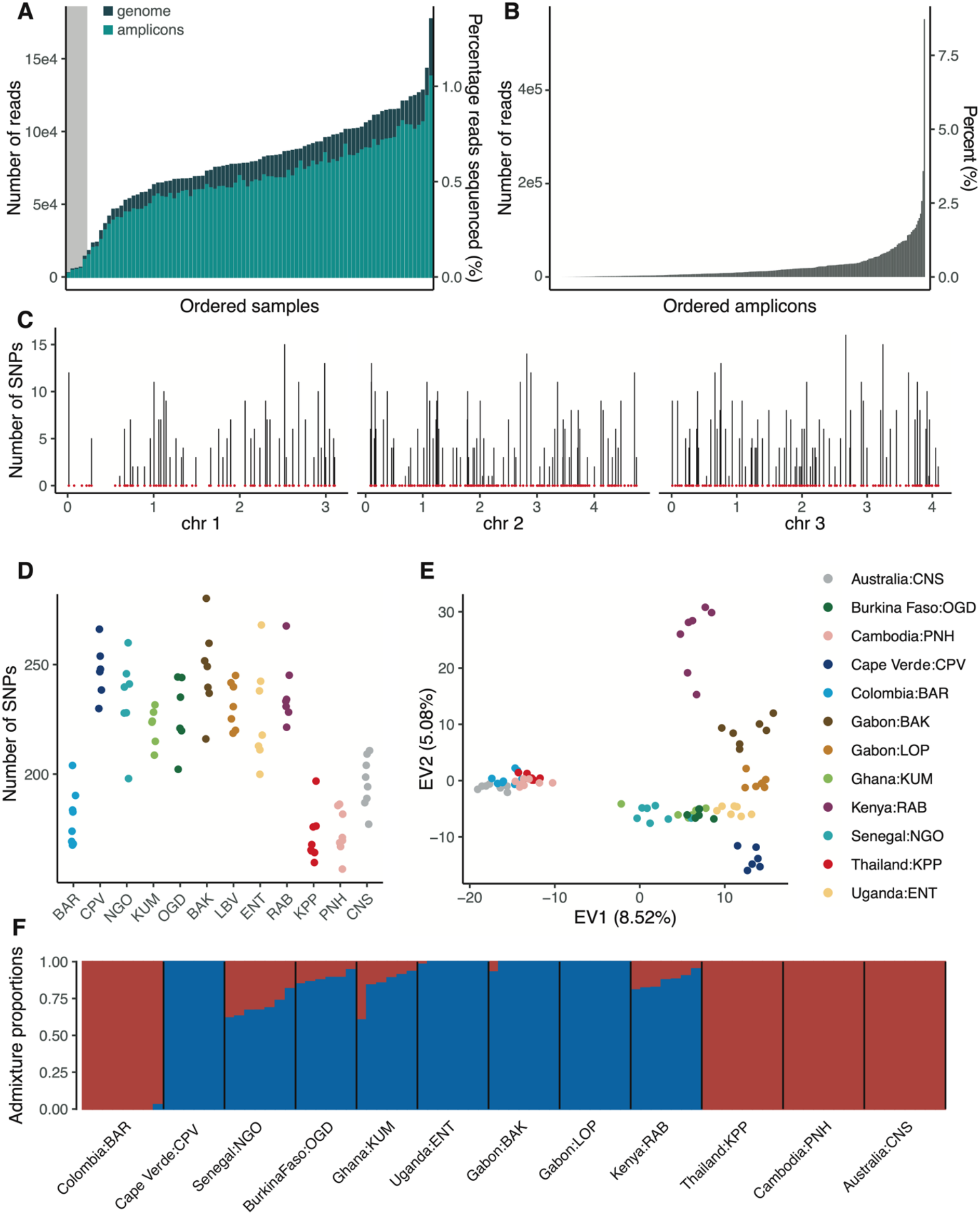
Genetic diversity and population structure of 12 laboratory colonies of *Ae. aegypti* determined by amplicon sequencing. Individual females from 12 colonies derived from field populations were analyzed after 7-29 generations (G) of laboratory colonization (BAR: Barranquilla, Colombia, G_19_; CPV: Praia, Cape Verde, G_7_; NGO: Ngoye, Senegal, G_17_; KUM: Kumasi, Ghana, G_15_; OGD: Ouagadougou, Burkina Faso, G_11_; BAK: Bakoumba, Gabon, G_26_; LOP: Lopé, Gabon, G_29_; ENT: Entebbe, Uganda, G_25_; RAB: Rabai, Kenya, G_18_; KPP: Kamphaeng Phet, Thailand, G_29_; PNH: Phnom Penh, Cambodia, G_26_; CNS: Cairns, Australia, G_23_). (**A**) Number of reads per sample mapped to the amplicon loci (light green) or the AaegL5 assembly (dark green), respectively. The grey shading indicates five samples that did not pass quality control and were excluded from subsequent analyses. (**B**) Number of reads uniquely mapping to each amplicon, ordered by increasing coverage. (**C**) Genomic distribution of amplicons (red dots) and number of SNPs identified at amplicon loci (vertical bars) along the three chromosomes (chr, shown in Mb) in the laboratory colonies (chr=chromosome). (**D**) Total number of SNPs identified per individual. (**E**) PCA of genetic diversity in the laboratory colonies at amplicon loci (EV=eigenvalue). (**F**) Barplot of admixture proportions per individual assuming two ancestry components (K=2, with *Aaa* ancestry shown in red and *Aaf* ancestry shown in blue).

We inferred the genetic ancestry composition in the laboratory colonies and found admixture patterns that broadly matched expectations based on their geographic origins (Figure 4F). Colonies originating from Africa showed a predominance of the generalist (*Aaf*) ancestry, with varying levels of admixture from the human-specialist (*Aaa*) ancestry, whereas non-African populations exhibited exclusive *Aaa* ancestry. Surprisingly, the colony originating from Cape Verde showed no signs of admixture, which was unexpected given the characteristics of the wild population it was derived from (Rose et al., 2022).

To assess the reproducibility of amplicon coverage across multiple sequencing runs, we prepared a new sequencing library with a subset of 75 individuals from the 12 laboratory colonies. Sequencing this new library produced between 2,660 and 45,006 reads per sample mapping to the amplicon loci, which displayed adequate quality and mapping statistics overall (Figure S1). Amplicon coverage was highly consistent between the two sequencing runs, as demonstrated by a strong and significant correlation in the percentage of reads mapped to each amplicon (R^2^=0.81, p<0.01; Figure S2). Our *in silico* analysis identified 60 primer pairs that were not specific to a single locus in the *Ae. aegypti* genome when allowing for a single mismatch. In both sequencing runs, amplicons with potential off-target loci did not show increased raw read coverage compared to the rest of the panel, however they displayed significant enrichment in multi-mapping reads (Figure S3). Of the 60 amplicons with putative off-target loci, 92% and 86% were covered by multi-mapped reads in each run, respectively, whereas these proportions were 1.2% and 0.8% for the rest of the amplicons. To avoid any bias due to reads derived from off-target loci, all multi-mapping reads were excluded from downstream analyses.

### Cost-effectiveness of the amplicon panel

To guide future use of the panel, we aimed to determine the minimum sequencing depth necessary to accurately genotype the targeted polymorphisms without unnecessary excess. Using data from the sequencing run that yielded about 8 million reads for 90 samples, we performed a rarefaction analysis by computationally resampling from 1 million to 20 million reads (Figure S4). We found that the proportion of genotyped amplicons (defined as those with at least 10× in depth of coverage) per sample increased asymptotically with the number of reads, with the curve plateauing at approximately 100,000 reads per sample. However, even at high sequencing depth, some variation in the proportion of genotyped amplicons persisted. This suggests that, for certain samples, there is insufficient template material to recover missing genotypes, and that further increasing sequencing depth will not resolve this limitation.

We also estimated the cost per sample of our amplicon sequencing strategy, based on the current price list at our institutions (Table S2). These cost estimates are obviously subject to change and may also depend on pack size, special deals, and local taxes. We calculated the cost of generating 100,000 reads per sample, which we identified as the optimal read coverage based on the rarefaction analysis, using an 8-million read flowcell on an Illumina MiSeq platform. We estimated that at Institut Pasteur in France the genotyping cost was 8.5 EUR per sample for library preparation and 13.6 EUR per sample overall, including the sequencing cost. At Yale University in the United States, the genotyping cost was approximately 10.1 USD per sample for library preparation and 27.8 USD per sample overall, including the sequencing cost. To help reduce the genotyping costs, we compared two commercial kits (Qiagen DNeasy Blood & Tissue and Molecular Research Center Inc. DNAzol DIRECT) and a low-cost, homemade protocol (called Pat-Roman) for DNA extraction. We compared the three DNA extraction methods based on the number of reads and SNPs genotyped for the same set of samples processed within the same library. Our analysis showed that the Pat-Roman protocol performed comparably to the Qiagen DNeasy Blood & Tissue kit, whereas the DNAzol DIRECT kit yielded poorer results (Figure S5). By implementing the Pat-Roman DNA extraction protocol and substituting plate normalization with manual normalization, we reduced the genotyping cost at Institut Pasteur to 4.6 EUR per sample for library preparation and 9.6 EUR per sample in total, including sequencing. At Yale University, the genotyping cost was reduced to 3.8 USD per sample for library preparation and 21.6 USD per sample in total, including sequencing.

## Discussion

In this study, we present a targeted amplicon sequencing strategy designed to enhance population genetic research on *Ae. aegypti*. This amplicon panel encompasses 291 loci throughout the *Ae. aegypti* genome and offers an accessible alternative to WGS, which is often prohibitive due to the large size and high repeat content of the *Ae. aegypti* genome. By focusing on selectively neutral loci, the panel mirrors the genetic diversity and population structure typically inferred from WGS data, thus providing a valuable tool for large-scale genotyping studies. The amplicon panel will complement existing reduced-representation approaches such as RAD-seq (Fontaine et al., 2017; Rasic et al., 2014; Rasic et al., 2015; Schmidt et al., 2018) and the commercially available *Ae. aegypti* SNP chip (Evans et al., 2015), without analytical challenges due to pervasive missing data (Arnold et al., 2013; Huang and Knowles, 2016) or ascertainment bias (Albrechtsen et al., 2010).

Our results demonstrate that the amplicon panel effectively distinguishes genetic diversity and structure within and among *Ae. aegypti* populations globally. The panel’s robustness is supported by the consistency of our findings with known genomic patterns, such as higher SNP diversity in African populations and separation of global populations in three distinct PCA clusters (Crawford et al., 2025; Lozada-Chavez et al., 2025). Furthermore, the panel’s ability to accurately estimate genetic ancestry components in naturally admixed populations highlights its utility in assessing admixture patterns in *Ae. aegypti*.

Our targeted amplicon sequencing panel provides a practical solution for genotyping *Ae. aegypti* at a fraction of the cost and resources required for WGS. We showed that the cost-effectiveness of the amplicon panel can be further improved by using a low-cost DNA extraction method and substituting plate normalization by manual normalization. While both modifications are more labor-intensive, they result in significant cost reductions, halving the cost of library preparation and reducing the total cost per sample by about 25%. Costs could be further reduced by using minimally invasive methods, such as incubating a gently squashed mosquito in lysis buffer (Korlevic et al., 2021; Makunin et al., 2022), then directly incorporating the crude lysate into PCR, avoiding the need for homogenization, complex buffers, or precipitation steps.

Due to the reduced-representation strategy of amplicon sequencing, there is a trade-off in genomic coverage resulting in fewer SNPs compared to WGS, which may impact the resolution of certain population genetic analyses. Our rarefaction analysis identified 100,000 reads per sample as the optimal coverage point, yet some variability in amplicon coverage remains, potentially due to variations in template material quality. Another limitation of the amplicon panel is the presence of multi-mapping reads, which must be excluded from downstream analyses. These multi-mapping reads presumably derive from off-target loci for a subset of 60 amplicons, which had lower uniquely-mapping read coverage compared to the rest of the panel.

In conclusion, our newly developed amplicon panel provides a cost-effective solution for genotyping *Ae. aegypti* on a large scale, especially in resource-limited settings. Future enhancements could focus on expanding the panel to include more loci for vector species identification and detection of bacterial symbionts such as *Wolbachia* (Batovska et al., 2018; Cannon et al., 2021; Lima-Cordon et al., 2025; Makunin et al., 2022). Additionally, these improvements could include screening for insecticide resistance (Campos et al., 2022; Collins et al., 2022; Fontaine et al., 2024; Lima-Cordon et al., 2025) and identification of large-scale chromosomal inversions using diagnostic SNPs (Liang et al., 2025). Our targeted amplicon sequencing strategy opens the door for extensive population surveillance and vector control studies, enhancing our understanding of genetic dynamics, which in turn can inform public health strategies.

## Acknowledgements

We thank Catherine Lallemand for assistance with mosquito rearing, Hervé Blanc for technical advice, all members of the Lambrechts lab for their insights, and Noah Rose for valuable feedback on the manuscript. We are grateful to Noah Rose, Carolyn McBride, Claudia Romero-Vivas, Silvânia da Veiga Leal, Massamba Sylla, Jewelna Akorli, Sampson Otoo, Athanase Badolo, Christophe Paupy, Diego Ayala, Davy Jiolle, John-Paul Mutebi, Joel Lutomiah, Rosemary Sang, Alongkot Ponlawat, Veasna Duong, and Gordana Rašić, for their role in establishing the *Ae. aegypti* colonies prior to this study.

## Funding

This work was funded by the French Government’s Investissement d’Avenir program, Laboratoire d’Excellence Integrative Biology of Emerging Infectious Diseases (grant ANR-10-LABX-62-IBEID to L.L.), and Agence Nationale de la Recherche (grant ANR-20-CE35-0002 to L.L.). S.N.R. and M.I.B. received support from a research-grant provided to Yale School of Public Health by the Ambrose Monell Foundation. Sequencing was performed at the Biomics core facility (C2RT, Institut Pasteur, Paris, France) supported by France Génomique (ANR-10-INBS-09) and IBISA, and at the Yale Center for Genome Analysis supported by the National Institute of General Medical Sciences of the National Institutes of Health under Award Number 1S10OD030363-01A1. The funders had no role in study design, data collection and analysis, decision to publish, or preparation of the manuscript.

## Competing interests

The authors declare that they have no competing interests.

## Authors contributions

Conceived and designed the study: J.D., S.N.R., L.L. Collected the data: A.L., L.M., P.M., M.I.B., I.D-B, P.K. Performed the analyses: J.D. Wrote the manuscript: J.D., L.L. Acquired the funding: S.N.R., L.L. All authors read and approved the final manuscript.

## Supplementary figures

**Figure S1:**
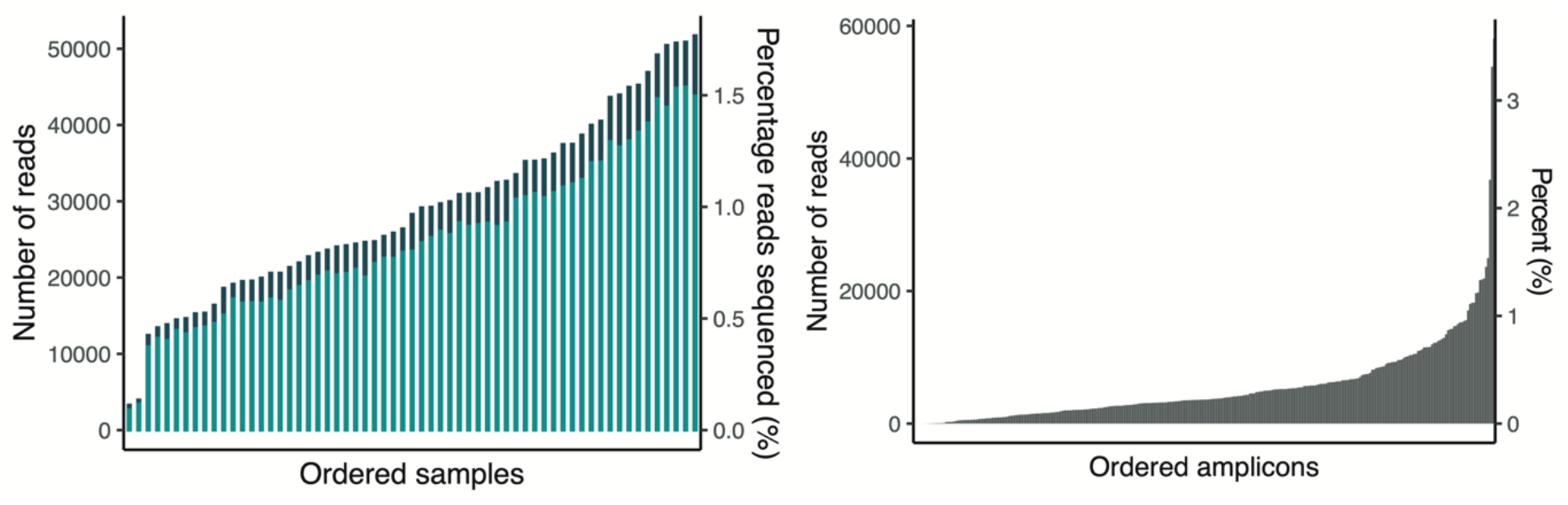
Mapping statistics of the technical replicate run. (**A**) Number of reads per sample mapped to the amplicon loci (light green) or the AaegL5 assembly (dark green), respectively. (**B**) Number of reads uniquely mapping to each amplicon, ordered by increasing coverage.

**Figure S2.**
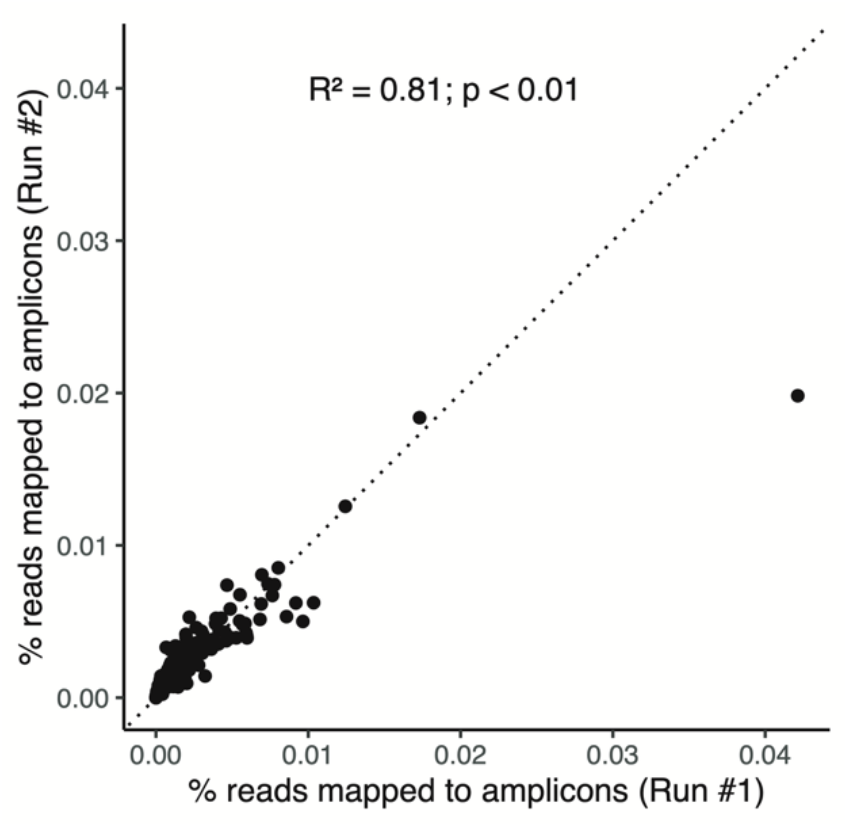
Amplicon sequencing reproducibility. Correlation between the percentage of reads mapped to each amplicon locus in the two replicate sequencing runs.

**Figure S3:**
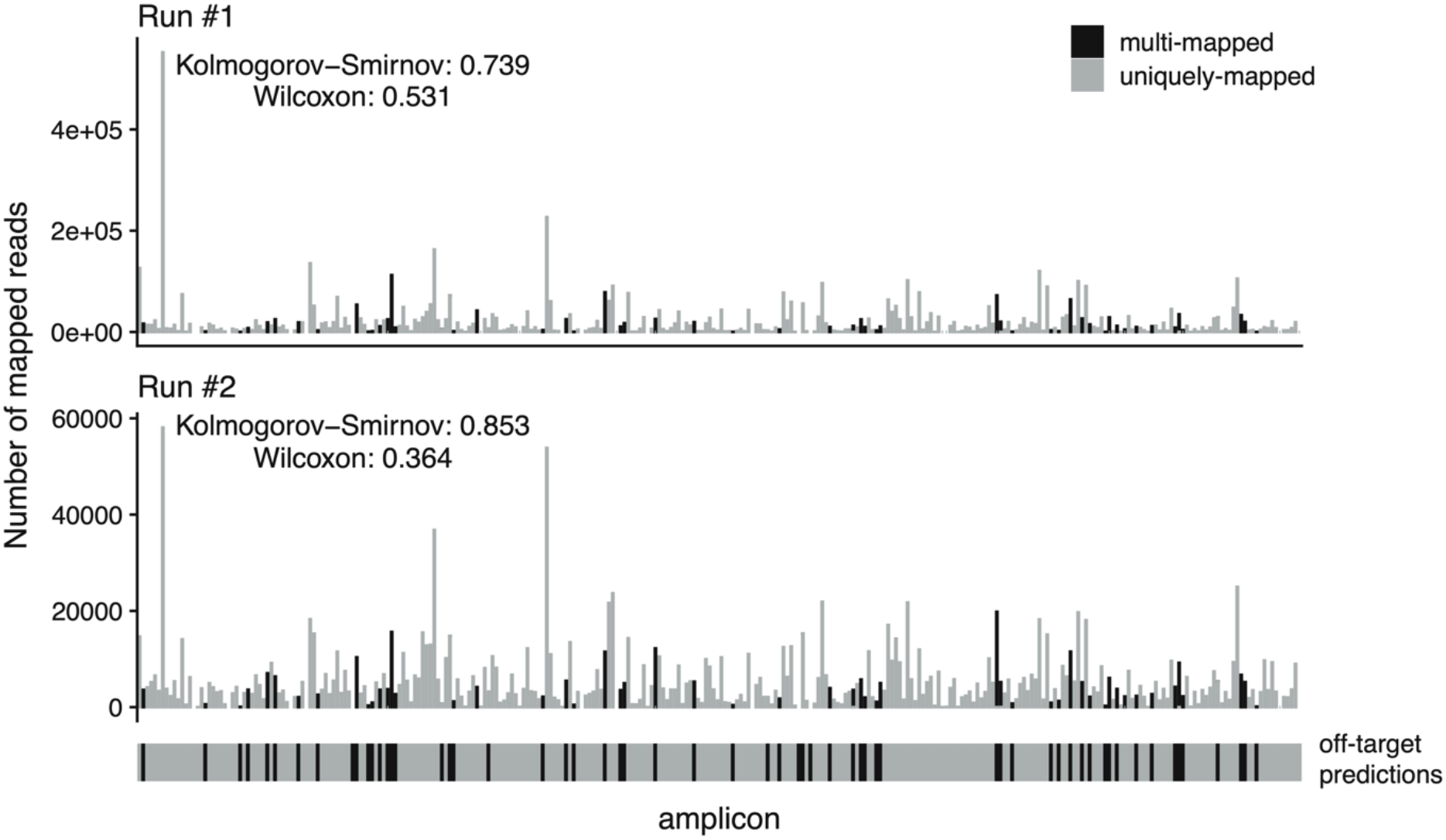
Visualization of amplicons with putative off-target amplification. The bottom panel shows the 291 amplicons, with vertical black lines indicating primer pairs that matched multiple sites in the genome. The top panel shows the number of reads mapped to each amplicon; gray bars represent uniquely mapped reads, while black lines represent multi-mapped reads. The Kolmogorov-Smirnov and Wilcoxon tests were conducted to assess whether there was a difference in the number of mapped reads between amplicons with predicted off-target loci and the other amplicons.

**Figure S4:**
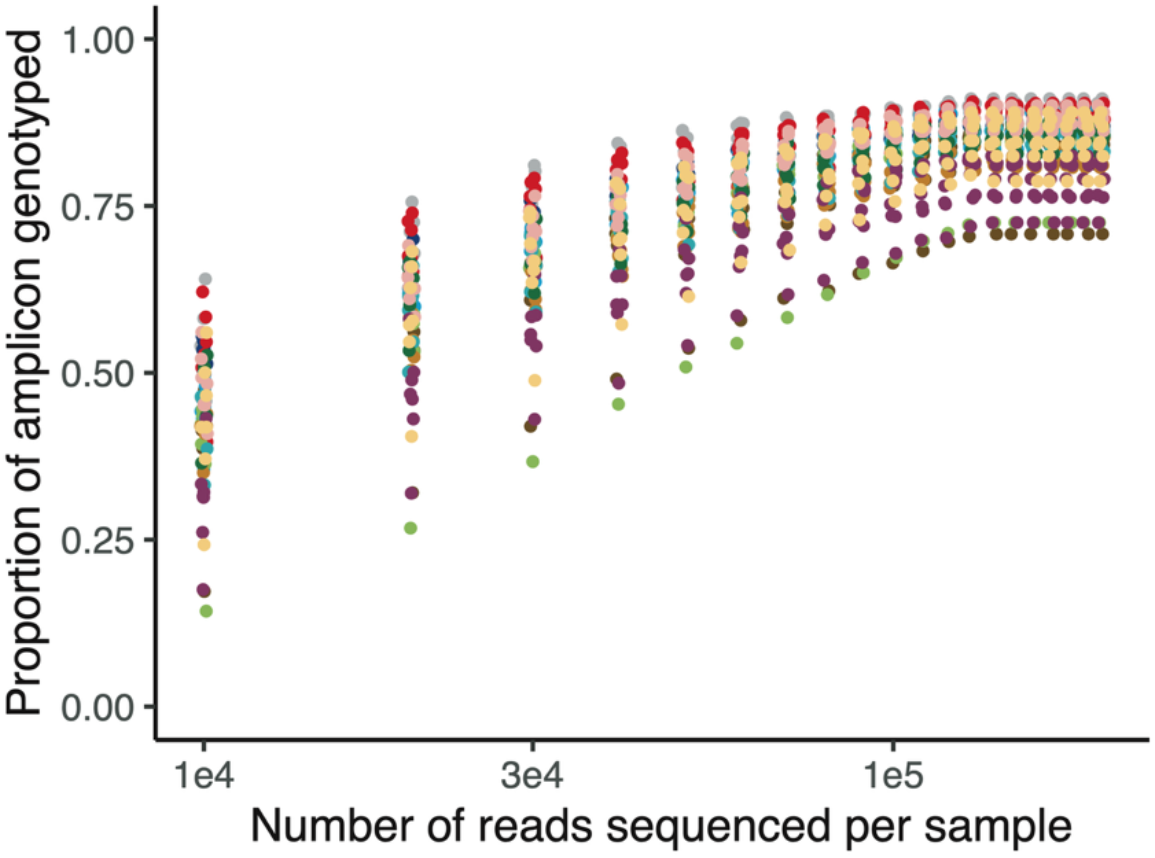
Rarefaction analysis of the relationship between the amplicon coverage and sequencing effort. The graph shows the proportion of amplicons genotyped (>10× coverage) as a function of the read counts per sample. This analysis was performed by computationally resampling from 1 to 20 million reads using data from a sequencing run that yielded about 8 million reads. Different colors represent individual samples.

**Figure S5.**
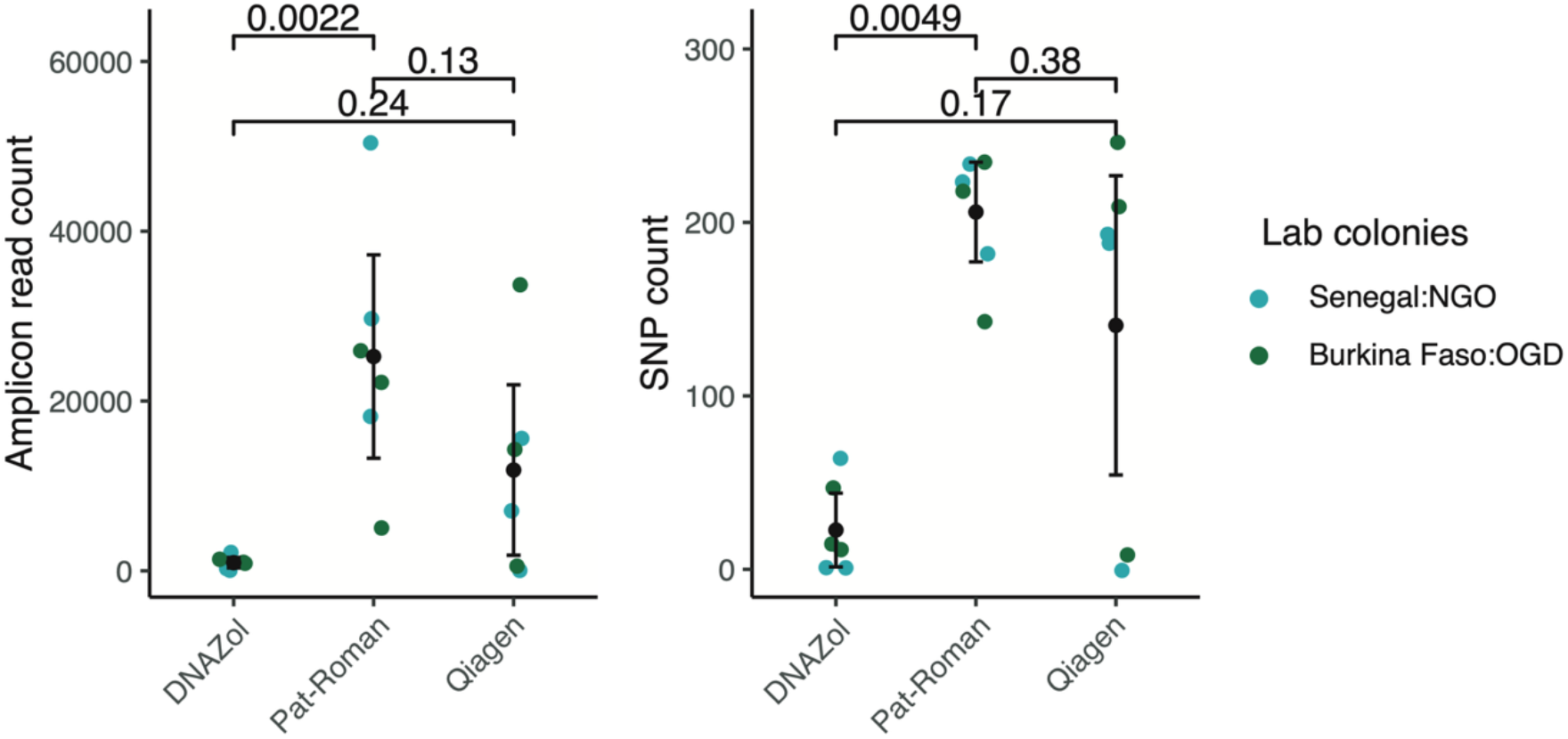
Comparison of three DNA extraction methods. Two commercial kits (Qiagen DNeasy Blood & Tissue and Molecular Research Center Inc. DNAzol DIRECT) and a low-cost homemade method (Pat-Roman) were compared based on the number of reads and SNPs genotyped from the same set of samples processed in the same library. The left panel shows the number of reads mapped to amplicon loci per sample, whereas the right panel displays the total number of SNPs identified. The data points are individual samples color-coded by colony, and the black vertical bar indicates the mean and 95% confidence interval. The p-values above the graphs were obtained using Wilcoxon test.

## Supplementary table legends

**Table S1: Amplicon loci and primers**. For each amplicon, the table provides the genomic positions, primer sequences, annealing temperature (Tm), and balancing factors.

**Table S2: Cost estimates**. The cost per sample was estimated based on the current price list at Institut Pasteur (France) and Yale University (United States). The number of samples per Illumina MiSeq Micro flowcell (n=80) was set to obtain 100,000 reads per sample, which we identified as the optimal read coverage based on the rarefaction analysis.

## Notes

### Competing Interest Statement

The authors have declared no competing interest.

